# Gene Regulatory Networks Mediate Pattern Scaling in Growing Tissues

**DOI:** 10.64898/2026.07.08.737218

**Authors:** Amy Bowen, Zena Hadjivasiliou

## Abstract

Developmental patterns can scale with size during growth, a phenomenon commonly attributed to morphogen scaling. Although patterning is orchestrated by gene regulatory networks (GRNs) activated by morphogens, how GRN dynamics interact with growth is not understood. We present a theoretical framework that integrates morphogen signalling, GRN dynamics, and tissue growth. We show that pattern scaling emerges from the interplay of GRN dynamics and growth, even in the absence of morphogen scaling. This relies on memory effects encoded in the GRNs, providing a cell-autonomous route to global scaling, and offering a general mechanism for size-invariant patterning beyond morphogen-based models.

---

During development, growth and patterning occur concomitantly, yet we understand little about their interplay. Pattern formation is often governed by the generation of gradients in signalling molecules known as morphogens, which effect gene expression via Gene Regulatory Networks (GRNs) [1–3] (Fig. 1 (a)). The interpretation of morphogens by GRNs influences patterning dynamics and plays a functional role in establishing precise and robust cell fate boundaries during development [4–8]. While theoretical advances have proposed models for the impact of growth and morphogenesis on morphogen gradients [9–12], we lack unifying frameworks that integrate growth with GRN dynamics. Recent measurements indicate that GRN dynamics and tissue growth can operate at similar timescales while the morphogen dynamics are significantly faster [13, 14]. This calls for theoretical approaches that explicitly couple growth with GRNs, and address their combined effect on tissue-level patterning.

In this work, we develop a theoretical framework that integrates morphogens, GRNs and growth to investigate how growth impacts patterning. We show that intrinsic multistability together with the non-negligible timescales of GRN dynamics relative to growth imbue cells with memory, shaping tissue-level patterning. This results in tissue-level pattern scaling, where the size of the regions defined by GRNs become proportional to the size of the growing tissue, despite a size-invariant morphogen gradient. Our results identify a scaling mechanism that does not require morphogen gradient scaling, but is instead driven by cell-autonomous GRN dynamics, shifting the focus of scaling from morphogens [15–19], to the GRN [20, 21].

**FIG. 1.**
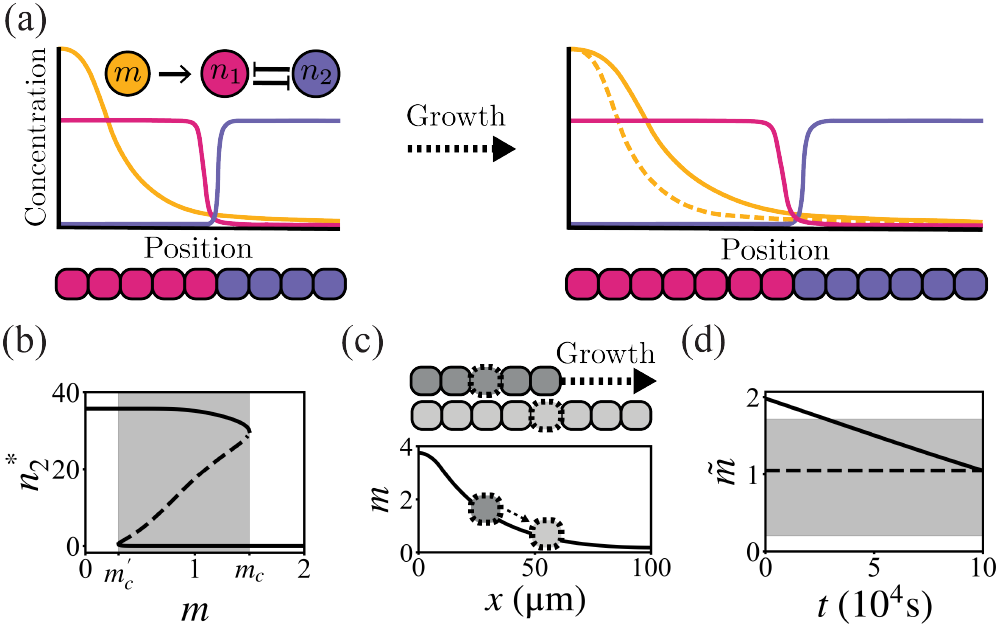
Morphogen and GRN patterning. (a) Morphogen induced toggle switch. Morphogen molecules form a concentration gradient (yellow) across position *x*. Gene concentrations (pink, purple) form spatially-dependent profiles. For larger tissues (right) patterns scale when gene profiles adapt to tissue length. The lengthscale of the morphogen may scale with tissue size (solid) or remain constant (dashed). (b) Bifurcation diagram showing the value of *n*_2_ at the fixed point, 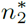 defined by critical values, against the morphogen concentration *m*. The bistable region (shaded), defined by critical values *m*_*c*_, 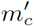, has 2 stable steady states (solid) and an unstable steady state (dashed). (c) Growth generates cellular flows, causing cells to move down the morphogen gradient *m*. (d) The morphogen signal a cell experiences through time, 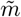 , may exit or enter the bistable region (shaded) for a growing tissue (solid) but stays static in a non-growing tissue (dashed).

## Patterning dynamics in growing tissues

We consider a growing two-dimensional system with morphogen concentration *m*(**x**, *t*) as a function of position **x** = (*x, y*) and time *t*. Morphogen dynamics depend on morphogen transport and degradation within the target tissue and local production within a source. In addition, growth affects the morphogen dynamics via advection and dilution [9, 11]. We further consider a mutually repressive two gene toggle switch network (Fig. 1(a)) where *n*_1_(**x**, *t*) and *n*_2_(**x**, *t*) are the gene concentrations for the nodes. The toggle switch is a common motif of GRNs that regulate patterning across developmental contexts and so a suitable minimal model for our analysis [1, 5, 8, 22–24]. Following [9], we consider morphogen and gene profiles that only vary along the *x* axis, and account for anisotropy of growth through *ε* = *∂*_*y*_*u*_*y*_*/∂*_*x*_*u*_*x*_ where *u*_*x*_ and *u*_*y*_ are the x- and y-components of the local cell velocity field **u**(**x**, *t*) and the growth rate is given by *g* = ∇ · **u**. The morphogen and gene dynamics are given by

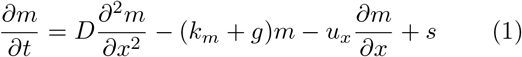

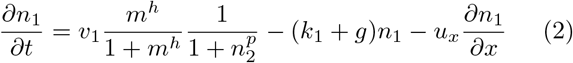

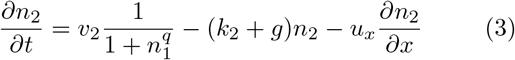

where *s* is the morphogen production term given by *s*(*x*) = *v*_*m*_ in a region 0 ≤ *x* ≤ *w* and *s*(*x*) = 0 elsewhere; *D* is the morphogen effective diffusion coefficient; *k*_*m*_, *k*_1_ and *k*_2_ are effective degradation rates and *v*_*m*_, *v*_1_ and *v*_2_ maximum production rates for normalized *m, n*_1_ and *n*_2_ respectively (Supplementary S1). The effects of gene activation or inhibition are expressed as Hill functions on the production terms, with Hill coefficients *h, p*, and *q* [1]. We assume reflective boundary conditions for the morphogen and genes at *x* = 0 and *x* = *L*(*t*). Equations 2 and 3 describe a bistable toggle switch where the production rate of *n*_1_ is modulated by the morphogen concentration *m*, which acts as a bifurcation parameter affecting the number and stability of attractors. At critical values of the morphogen *m*_*c*_, *m*′_*c*_, with corresponding positions *x*_*c*_, *x*′_*c*_, the system undergoes a saddle-node bifurcation, demarcating a bistable region in space where, for a given signal *m*, the initial values of *n*_1_ and *n*_2_ determine which basin of attraction the system settles in (Fig. 1(b)).

We consider a growth rate that is constant in space and time. The position of individual cells *ψ* Therefore evolves exponentially in time,

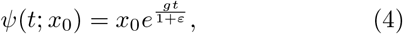

where *x*_0_ is the initial cell position. The size of the tissue thus evolves as 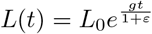 for an initial tissue length *L*_0_. The velocity field is linear in space for spatially uniform and constant growth, given by *u*_*x*_(*x, t*) = *gx/*(1 + *ε*) and is effectively time independent.

Growth generates movement of cells through advection, which affects the morphogen signal a cell experiences in time (Fig. 1(c)). We assume that morphogen turnover is faster than growth [13, 14, 25]. In this limit, morphogen degradation dominates over growth effects, *k*_*m*_ ≫ *g*, the morphogen gradient can be approximated by its steady state solution (Supplementary S1). In the co-moving frame of a cell with position *ψ*(*t*; *x*_0_) outside the source, the morphogen signal experienced is given by,

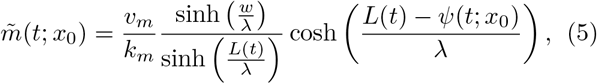

where 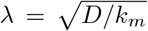 the lengthscale of the morphogen gradient. The dynamics of 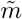 can push cells in or out of the bistable region (Fig. 1(d)), at a rate dependent on the morphogen amplitude *v*_*m*_*/k*_*m*_, decay length *λ*, and growth rate *g*. This differs from the case where the tissue does not undergo growth, and a cell in a given position experiences a constant 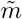 (Fig. 1(d)).

## Scaling emerges due to bistability

In the limit *k*_*m*_ ≫ *g*, the morphogen gradient is unaffected by tissue growth (Fig. 2(a)). To extract the effects of growth we explore two cases: patterning in non-growing, static tissues of different sizes and patterning in growing tissues. For static tissues, we expect no pattern scaling with tissue size since the morphogen that directs patterning is constant for each cell and independent of tissue size. Given that advection causes cells to experience dynamic morphogen signals (Fig. 1(c,d)), we explore how growth affects patterning.

**FIG. 2.**
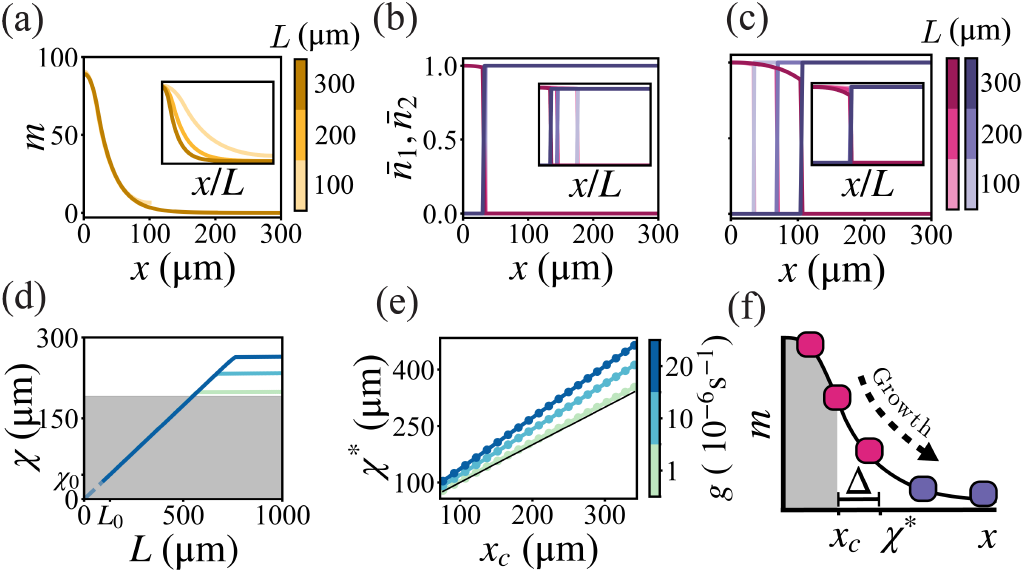
Scaling is mediated by GRN multistability. (a)-(c) Concentration versus position for tissue lengths *L* = [100, 200, 300]µm (colourbars); insets in relative coordinates *x/L*. 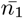 and *n*_2_ the gene concentrations normalised by their maximum amplitudes. (a) Morphogen concentration *m* versus position *x* for different *L*. (b) *n*_1_ and *n*_2_ concentration profiles for different *L* for a static, non-growing tissue. (c) *n*_1_ and *n*_2_ concentration profiles for different tissue lengths for a growing tissue. (d) Boundary position *χ* against tissue length *L* for growing tissues, with different growth rates (colourbar). Pattern boundary extends beyond the bistable region (grey, shaded). (e) The maximum boundary position *χ*^*^ plotted against the position of the edge of the bistable region *x*_*c*_, for different growth rates (colourbar). *x*_*c*_ was varied by changing the morphogen diffusion coefficient *D*. Black line shows *χ*^*^ = *x*_*c*_. (f) Schematic of cells advected down the morphogen gradient. Cells retain a high *n*_1_ concentration beyond the edge of the bistable region *x*_*c*_ (shaded grey) by an extension Δ, which together define the maximum boundary position *χ*^*^.

We quantify the patterning dynamics by finding find the position of the boundary between high *n*_1_ and high *n*_2_ regions, which we term *χ*(*t*) (Supplementary S2, Fig. S1). For static tissues, we find the *n*_1_ and *n*_2_ profiles are unaffected by *L*, as expected (Fig. 2(b)) and the boundary position is constant at *χ*_0_. When we instead initiate the system at tissue length *L*_0_ and introduce uniform growth, we find the *n*_1_ and *n*_2_ profiles in absolute coordinates change over time, while the relative profiles are invariant (Fig. 2(c)), and the position of the boundary *χ* scales linearly with the tissue length *L* (Fig. 2(d)). Scaling fails beyond a maximum tissue size which corresponds to a maximum value of *χ*(*t*) which we term *χ*^*^. The value of *χ*^*^ depends on the position of the edge of the bistable region *x*_*c*_, and on the growth rate *g*, suggesting that both GRN bistability and growth of the domain underlie the emergent scaling. Accordingly, for systems that do not exhibit multistability, we find no scaling for small growth rates (Fig. S2), while for other network topologies with multistability, such as the self-activating node, the relationship between scaling, multistability and growth holds (Fig. S3).

The relationship between the edge of the bistable region *x*_*c*_ and the boundary position *χ* can be understood as follows. In a non-growing tissue, the morphogen signal experienced by a cell 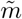 is constant in time, since cell position is static and the morphogen is at steady state. Therefore, the boundary position for the static case, *χ*_0_, occurs at a threshold morphogen concentration defined by the network parameters and the initial conditions (Fig. S4) and is independent of tissue length. In a growing tissue, however, the morphogen signal experienced 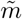 decays over time (Fig. 1(d)). A cell that begins close to the morphogen source (*x*_0_ *< χ*_0_), initially specifies to a high *n*_1_ fate, yet as advection pushes the cell down the morphogen gradient (Fig. 1(c)), the cell travels through the bistable region. So long as a cell remains in the bistable region, it remains at the high *n*_1_ attractor. This results in the boundary travelling at the advection velocity, as the cell at the boundary retains memory of its previous higher morphogen environment. For uniform growth, this maintains the relative position of the boundary in *x/L* space, thus scaling the system *χ*_0_*/L*_0_ = *χ*(*t*)*/L*(*t*). We have also explored how initial conditions and the structure of the *n*_1_ and *n*_2_ bifurcation diagram impact scaling, and found that these have quantitative but not qualitative effects (Figs. S4,5).

A logical consequence of the arguments above is that scaling will cease once the boundary position surpasses the bistable region. For small growth rates, *g* → 0, *χ*^*^ tends to the position marking the edge of the bistable region *x*_*c*_ (Fig. 2(d,e)), consistent with this prediction. However, when we increase the growth rate, *χ*^*^ extends beyond *x*_*c*_, suggesting an additional mechanism of scaling. We define Δ as the extension to the maximum fate boundary beyond the edge of the bistable region, *χ*^*^ = *x*_*c*_ + Δ (Fig. 2(f)). Larger growth rates lead to larger Δ (Fig. 2(d,e)) suggesting this mechanism of scaling is growth-dependent.

## Relative timescales of GRN and growth expand scaling

The dependence of the maximum boundary position for scaling, *χ*^*^, on the growth rate results from the combined effects of advection and the dynamics of signalling. Consider a cell that initially experiences a high morphogen signal, and specifies to a high *n*_1_ state. When the cell is advected to the edge of the bistable region (Fig. 3(a)), the high *n*_1_ attractor disappears, leaving only the high *n*_2_ attractor (Fig. 3(b)). The cell respecifies to the high *n*_2_ attractor at a rate affected by both the network parameters and the rate at which the morphogen decays in the frame of the cell. The decay from a high *n*_1_ state to a high *n*_2_ state is associated with a timescale we define as *τ* (Fig. 3(b)). The extension to scaling beyond the bistable region, Δ, corresponds to the distance traversed by a cell due to advection in the time it takes for cells to respecify from a low to a high *n*_2_ state outside the bistable region (Fig. 3(a,b)). We numerically estimate *τ* as the time it takes upon exiting the bistable region to reach the threshold *n*_2_ concentration that defines the boundary *χ* (Fig. 3(c)). It follows that Δ is given by,

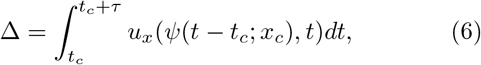

where we integrate the velocity over time of a cell that exits the bistable region at *t* = *t*_*c*_ and follows a trajectory *ψ*(*t t*_*c*_; − *x*_*c*_) (Fig. S1).

**FIG. 3.**
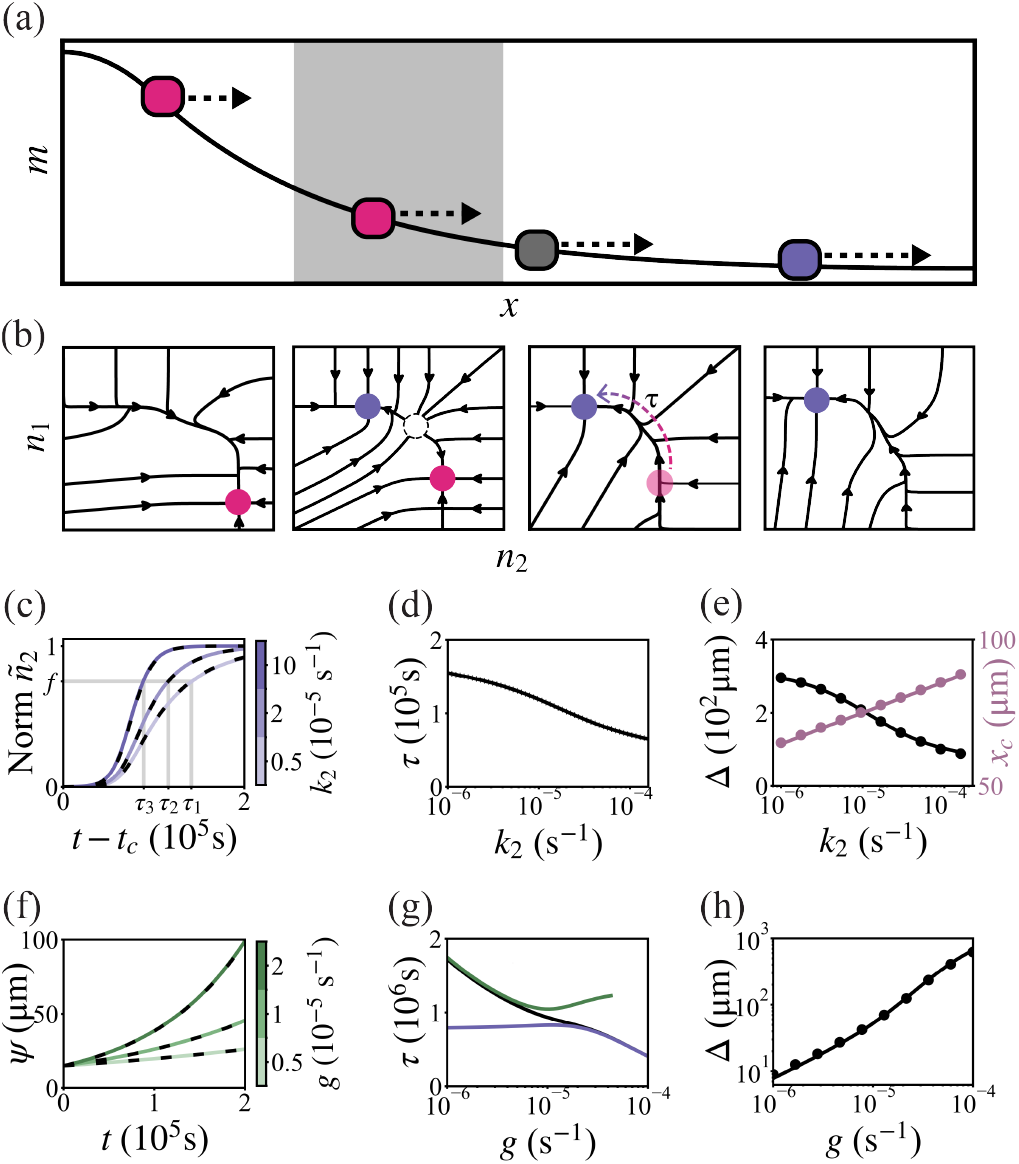
Slow relaxation of the GRN mediates scaling. (a) A cell travelling via advection (arrows) through the morphogen gradient *m* across position *x*, from the high *n*_1_ monostable region to the bistable region (shaded) and high *n*_2_ monostable region. Arrows indicate the direction and magnitute of advection. (b) *n*_1_ *− n*_2_ phase portraits associated with cells in for different values of morphogen *m* as they are advected through the tissue. (c) The normalised *n*_2_ concentration of a given cell, *ñ*_2_, versus time since leaving the bistable region, *t − t*_*c*_, for varying *k*_2_ (colourbar), from numerical simulations (solid) and analytical expressions of 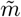 input into a non spatial solver for *ñ*_1_, *ñ*_2_ (dashed, black). The respecification times for the different *k*_2_ values, *τ* , are defined when the normalised *ñ*_2_ reaches a fraction *f* . (d) Respecification time *τ* against degradation rate *k*_2_. (e) Extension to scaling Δ ( black) and position of the edge of the bistable region *x*_*c*_ (purple) against degradation rate *k*_2_. 1D numerical solutions for Δ (black, circles) and analytical solutions (black line) obtained from Equation 6 with numerical input of *τ* . (f) Cell position *ψ* against time *t* for different growth rates, *g* (colourbar), from numerical simulations (solid) and analytical solutions from Equation 4 (dashed, black). (g) Respecification time *τ* against the growth rate *g* (black). Simulations without dilution (green) and at the limit of fast advection (purple). (h) Extension to scaling Δ against growth rate *g* from 1D numerical solutions (circles) analytical solutions (line) obtained from Equation 6 with numerical value of *τ* .

Considering advection at the edge of the bistable region results in the following expression (Fig. S6),

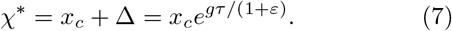

To quantitatively determine how the growth rate and network dynamics each affect the extension to scaling, we explore how *τ* and Δ depend on advection, dilution and the network dynamics. The timescale *τ* depends on how quickly the morphogen signal is changing, and how rapidly the network processes the changing signal. Increasing degradation rates speeds up the dynamics (Fig. 3(c)). Accordingly, *τ* decreases when we increase *k*_2_, the degradation rate of *n*_2_ (Fig. 3(d)). Varying *k*_2_ has the additional effect of changing the position of the bistable region, *x*_*c*_, as the critical morphogen concentrations depend on the underlying network parameters (Fig. 3(e)). An increase in *x*_*c*_ leads to an increase in the advection velocity at the edge of the bistable region, which increases Δ, Equation 7. A decrease in *τ* , however, means that the cell has less time to advect before respecifying, acting to decrease Δ: this effect dominates when the degradation rate *k*_2_ is limiting the GRN timescale.

We next consider how *τ* and Δ are influenced by growth. The spatial position of cells, *ψ*, increases exponentially over time (Equation 4, Fig. 3(f)). For higher growth rates, cells move further due to advection over the same time period, increasing Δ. However, the *n*_1_ and *n*_2_ dynamics are also influenced by advection which affects 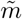 , and by dilution which introduces an effective degradation rate, thereby influencing the respecification time *τ* (Fig. 3(g)). We decompose these effects by numerically calculating *τ* for two limiting cases: first, a system without the dilution term; and second, a system at the limit of fast advection, corresponding to a signal 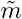 that decays rapidly upon exiting the bistable region (Fig. 3(g)). The effect of dilution acting as an effective degradation rate dominates for large growth *g* ≫ *k*_1_, *k*_2_ . For small growth, advection dominates the decrease in *τ* by speeding up the decay of 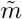 . Advection also acts to increase Δ by increasing the velocity of cells. Altogether, the increase in advection velocity dominates over the decrease in *τ* , leading Δ to increase with growth rate (Fig. 3(h)).

In summary, scaling can extend beyond the bistable region by a length Δ, which emerges due to the timescale of network relaxation relative to the cell movement due to advection. This effect is independent of multistability and persists in regimes where the toggle switch is monostable (Fig. S2). The constituent processes of advection and network dynamics are influenced antagonistically by the network parameters and growth rate, highlighting the non-trivial interaction of network and growth dynamics during patterning (Fig. S7).

## Morphogen source growth extends scaling

In biological systems, tissue growth is often accompanied by increase in the size of the morphogen source [25–27], which affects the morphogen profile, and therefore changes the position of the edge of the bistable region (Fig. 4(a)). We ask how source growth interacts with scaling mediated by GRN dynamics.

**FIG. 4.**
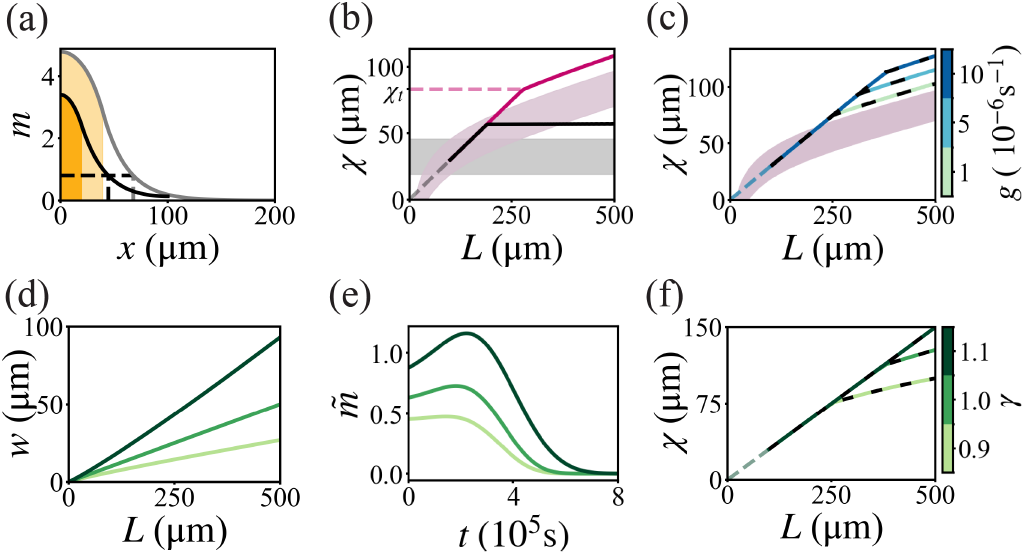
Source growth extends scaling. (a) Morphogen concentration *m* versus position *x* for a small (dark shaded, black line) and larger (light shaded, grey line) source, and respective threshold positions (dashed line). (b) Boundary position *χ* versus tissue length *L*, without (dashed, grey) and linear (solid, purple) source growth, and respective bistable regions (shaded). (c) Boundary position *χ* versus tissue length *L* for different growth rates, *g* (blue colourbar) for *γ* = 1, numerical simulations (solid) and analytical solutions (dashed). (d) Source width *w* versus tissue length *L* for different *γ* (green colourbar). (e) Morphogen signal experienced by a cell 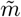 at relative position *x/L* = 0.3 over time for different *γ* (green colourbar). (f) Boundary position *χ* versus tissue length *L* for source growth modes *γ* (green colourbar), numerical simulations (solid) and analytical solutions (dashed).

We investigate source width *w* that scales with tissue length, *L, w* = *βL*^*γ*^. For linear source growth, *γ* = 1, the system initially behaves exactly as for a static source: *χ* increases proportionally to *L* leading to pattern scaling (Fig. 4(b)). For a static source, scaling ceases for *χ > χ*^*^, however, with source growth, *χ* continues to increase, tracing the evolution of the edge of the bistable region, *x*_*c*_(*t*) (Fig. 4(b)). We recapitulate these dynamics by accounting for the dynamics of the bistable region, *x*_*c*_(*t*), which becomes time-dependent as the morphogen source grows, and define a piecewise function for *χ*,

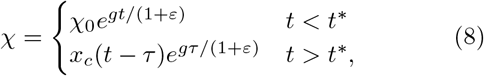

where *t*^*^ is defined by 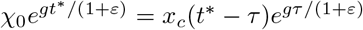 (Supplementary S9). As the bistable region expands, cells retain their high *n*_1_ fate further into the tissue, extending the boundary. An absolute upper limit for the patterning length-scale no longer exists. The dynamics of *χ* are biphasic: initially *χ* evolves due to advection, until it reaches a limit *χ*_*t*_ imposed by the instantaneous position of the edge of the bistable region plus the extension Δ emerging from GRN dynamics. The value of *χ*_*t*_ depends on the magnitude of advection compared to source growth, and increases with the growth rate (Fig. 4(c)). Beyond *χ*_*t*_, *χ* follows *x*_*c*_(*t*) +Δ, tracing the moving edge of the bistable region. Similar dynamics are present in slow-growing, monostable cases, where scaling is independent of GRN dynamics and arises due to source growth moving the morphogen concentration threshold defining the transition from the *n*_1_ to the *n*_2_ state (Supplementary S9).

We next explored sublinear and superlinear growth of the source width *w* with respect to tissue length *L*, by varying *γ* (Fig. 4(d)). Varying *γ* changes the dynamics of the morphogen signal experienced by a cell, 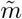 (Fig. 4(e)), by affecting the source width dynamics *w*(*t*). For larger values of *γ*, the increase in source width *w* dominates over the effects of increasing cell position *ψ*, causing 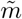 to reach a local maximum (Fig. 4(e)). This corresponds to the movement of the bistable region overtaking cell advection, and cells reach the edge of the bistable region *m*_*c*_ at a later time for increasing *γ*.

Although the initial evolution of *χ* is independent of source growth, at later times *χ* depends on the dynamics of the bistable region dominated by the source growth (Fig. 4(c)). As *γ* increases, the rate of increase of *x*_*c*_ with *L* increases, impacting the value of *χ*_*t*_ which defines the position where *χ* transitions from being advection dominated to source growth dominated. By considering the points at which the advection-driven scaled boundary exits the bistable region, constraints on *γ* can be found that ensure *χ* remains within the bistable region by virtue of the source growth (Fig. S8), increasing the value of *χ*_*t*_ and maintaining the same scaling slope for larger *L* (Fig. 4(f)).

In summary, the effect of source growth can be understood via its translation of the bistable region. As in the static source case, GRN relaxation adds a buffer zone of magnitude Δ around the bistable region that enables scaling to persist.

## Conclusion

We have developed a theoretical framework for modelling the effects of growth on patterning that explicitly captures the interaction of GRN dynamics with advection and dilution. Using this framework, we propose a mechanism for scaling in growing tissues whereby multistability and relaxation dynamics at the level of the GRN lead to pattern scaling at the tissue level. This demonstrates a mechanism for pattern scaling that does not emerge from scaling of the static positional information imparted by a morphogen gradient, but instead, utilises the dynamic and non-linear processing of the morphogen by GRNs.

Representing GRNs as dynamical systems recapitulates the dynamics of cell fate decisions [2, 7, 8, 28– 31]. Notably, bistability and hysteresis are common properties of nonlinear dynamical systems [32, 33] that have been evidenced in diverse developmental contexts [1, 8, 34, 35]. Growth naturally generates a time dependent morphogen input so that the GRN state of a cell may not map to the morphogen concentration at its current position, but instead retains information about the distance travelled due to advection. By explicitly considering the effects of growth on the morphogen input cells receive, we propose a mechanism by which the memory imbued by hysteresis in the GRN results in pattern scaling. Scaling in our analysis emerges from multistability, thus, could hold for networks that exhibit multistability or other types of bifurcations regardless of the exact structure of the network [36]. While we focus on growth, hysteresis and multistability can take effect in other scenarios in which the signal received is time-dependent, including morphogenetic flows [12, 37], cell movements [38, 39], noise at the level of the morphogen gradient [40–42], or out of steady state dynamics[43, 44].

Even in the absence of multistability, GRN dynamics have an effect on pattern scaling during growth. Experimental quantifications indicate that GRN dynamics and growth can operate at timescales of similar order of magnitude [13, 14]. In such cases, growth can impact GRN dynamics, affecting the time- and length-scales of tissue-level patterning. This result emphasises the impact of GRN dynamics on tissue level properties over relevant timescales of developmental events, and highlights the limitations of steady state assumptions, [44, 45].

In many developmental contexts, tissue growth is accompanied by growth of the morphogen source, which has been proposed to mediate scaling [26, 46]. We demonstrate how source growth alone can contribute to scaling and how source growth can interact with GRN memory effects to extend scaling synergistically.

In conclusion, this work highlights the need to integrate growth and cell movements when interrogating spatial and temporal dynamics of patterning in growing or, more broadly, dynamic environments. Considering the dynamics of cell position becomes particularly important when tissue flows or growth operate at timescales comparable to the timescales of GRN relaxation, and can drive emergent properties such as tissue-level pattern scaling.

## ACKNOWLEDGMENTS

We are grateful to James Briscoe, Jake Cornwall-Scoones, Lewis Mosby, Ruben Perez-Carrasco and Timothy Saunders for feedback on the manuscript and members of the Hadjivasiliou Lab for discussions . This work was supported by the Francis Crick Institute, which receives its core funding from Cancer Research UK, the UK Medical Research Council, and Wellcome Trust.

