## Supplementary Text for "Gene Regulatory Networks Mediate Pattern Scaling in Growing Tissues"

### Supplementary Material for: Gene Regulatory Networks Mediate Pattern Scaling in Growing Tissues

Amy Bowen[1,2,3] and Zena Hadjivasiliou[1,2,3,*]
[1] *Mathematical and Physical Biology Laboratory, The Francis Crick Institute, London, UK*
[2] *Department of Physics and Astronomy, University College London, London, UK*
[3] *London Centre for Nanotechnology, London, UK*

In this document, we provide details on the derivations presented in the main text and methods of pattern quantification. We provide additional results for scaling in a toggle switch within the monostable regime, and in a self-activating node system. We detail the effects of initial conditions and the shape of the bifurcation diagrams, specifically fixed point amplitude decay, on scaling. We show the full equation for boundary position dynamics and demonstrate the agreement between this equation and numerical simulations. We provide additional quantifications of the effect of GRN and growth parameters on the respecification time of GRNs in growing tissues. Finally, we provide derivations for equations regarding source growth and additional results for scaling mediated by source growth in combination with GRN dynamics.

#### S1. NON-DIMENSIONALISATION OF MORPHOGEN AND GRN EQUATIONS

We describe the morphogen concentration $M(\mathbf{x}, t)$ at position $\mathbf{x} = (x, y)$ and time $t$ on a growing domain of area $A(t)$ by a reaction-diffusion equation, where $D$ is the 2D diffusion coefficient, $k_M$ is the degradation constant, $u$ is the local flow velocity, $g$ is the growth, and $V_M$ is the morphogen production term. Equation S1 is a Partial Differential Equation describing the evolution of $M$ [1, 2],

$$\frac{\partial M}{\partial t} = D\nabla^2 M - k_M M - \mathbf{u}\cdot\nabla M - gM + S(\mathbf{x}). \quad \text{(S1)}$$

We consider morphogen profiles that only vary along the $x$-axis, allowing us to reduce the system to 1 dimension [2],

$$\frac{\partial M}{\partial t} = D\frac{\partial^2 M}{\partial x^2} - k_M M - u_x\frac{\partial M}{\partial x} - gM + S(x), \quad \text{(S2)}$$

where $u_x = \int_0^x g/(1+\varepsilon)dx'$, the $x$-component of the velocity that obeys $g = \nabla\cdot\mathbf{u}$. The anistropy parameter is defined by $\varepsilon = \partial_y u_u/\partial_x u_x$. The source term $S(x) = v_m$ in a region $0 \le x \le w$ and $S(x) = 0$ elsewhere.

The 1D equations for the dimensional gene concentrations $N_1(x,t)$ and $N_2(x,t)$ are as follows,

$$\frac{\partial N_1}{\partial t} = \frac{V_1(M/M^*)^h}{1+(M/M^*)^h}\frac{1}{1+(N_2/N_2^*)^p} - (k_1+g)N_1 - u_x\frac{\partial N_1}{\partial x} \quad \text{(S3)}$$

$$\frac{\partial N_2}{\partial t} = \frac{V_2}{1+(N_1/N_1^*)^q} - (k_2+g)N_2 - u_x\frac{\partial N_2}{\partial x}, \quad \text{(S4)}$$

where $M^*$, $N_1^*$ and $N_2^*$ are the half maximum concentrations for the respective Hill functions activating or inhibiting production, and $V_1$ and $V_2$ are production rates of $N_1$ and $N_2$ respectively.

We introduce the non-dimensional parameters $m = M/M^*, n_1 = N_1/N_1^*$ and $n_2 = N_2/N_2^*$, and scaled production rates $v_m = V_M/M^*$, $v_1 = V_1/N_1^*, v_2 = V_2/N_2^*$. Equations S2, S3 and S4 reduce to Equations 1, 2 and 3 in the main text.

#### S2. STEADY STATE LIMIT FOR MORPHOGEN DYNAMICS

The steady state solution to Equation 1 in the main text for a non-growing system, $g = 0$, with reflective boundary conditions is given by

$$m^{SS}(x) = \frac{v_m}{k_m}\begin{cases} 1 - \frac{\sinh\frac{L-w}{\lambda}}{\sinh\frac{L}{\lambda}}\cosh\frac{x}{\lambda} & x < w \\ \frac{\sinh\frac{w}{\lambda}}{\sinh\frac{L}{\lambda}}\cosh\frac{L-x}{\lambda} & x > w \end{cases} \quad \text{(S5)}$$

where $\lambda = \sqrt{\frac{D}{k_m}}$. We focus on the regime where $g \ll k_m$ where the steady state solution is a good approximation.

In the limit where the morphogen decay length is smaller than the tissue length, $\lambda \ll L$, Equation S5 simplifies to

$$m^{SS}(x) = \frac{v_m}{k_m}\begin{cases} 1 - e^{-w(t)/\lambda}\cosh\frac{x}{\lambda} & x < w(t) \\ \sinh\frac{w(t)}{\lambda}e^{-x/\lambda} & x > w(t). \end{cases} \quad \text{(S6)}$$

By rearranging Equation S5, one can find the position $x_c = x(m_c)$ that corresponds to the morphogen value $m_c$,

$$x(m_c) = \begin{cases} \lambda\cosh^{-1}\left(1 - \frac{m_c k_m}{v_m}\frac{\sinh(\frac{L-w}{\lambda})}{\sinh(\frac{L}{\lambda})}\right) & m_c < m_w \\ L - \lambda\cosh^{-1}\left(\frac{m_c k_m}{v_m}\frac{\sinh(\frac{L}{\lambda})}{\sinh(\frac{w}{\lambda})}\right) & m_c > m_w, \end{cases} \quad \text{(S7)}$$

where $m_w$ is the morphogen concentration at the edge of the source, $m_w = \frac{v_m}{k_m}\frac{\sinh\frac{w}{\lambda}}{\sinh\frac{L}{\lambda}}\cosh\frac{L-w}{\lambda}$.

In the limit $\lambda \ll L$ the expression above reduces to,

$$x(m_c) = \begin{cases} \lambda\cosh^{-1}\left(e^{w/\lambda}(1 - \frac{m_c k_m}{v_m})\right) & m_c < m_w \\ \lambda\ln\left(\frac{v_m\sinh(\frac{w}{\lambda})}{m_c k_m}\right) & m_c > m_w, \end{cases} \quad \text{(S8)}$$

where the morphogen concentration at the edge of the source in this limit is $m_w = \frac{v_m}{k_m} \sinh \frac{w}{\lambda} e^{-w/\lambda}$.

Equation S7 or S8 can be used to find the positions that demarcate the bistable region $x_c$ and $x_c'$, by substituting in $m_c$ as the critical morphogen concentrations for bistability. The critical morphogen concentrations can be found in implicit form for the bistable toggle switch [3].

Under the steady state assumption for the morphogen, the morphogen concentration experienced by a cell as it follows a position trajectory $\psi(t;x_0)$, which we denote by $\tilde{m}$, can by found from Equation S5 by transforming coordinates from the Eulerian frame in fixed space coordinate $x$ to the Lagrangian frame with co-moving coordinate $\psi$. In the main text, Equation 5 shows $\tilde{m}$ in the case where the cell is outside the morphogen source for all times, $\psi(t) > w(t)$. The general case for a time-dependent source $w(t)$ is given by,

$$\tilde{m}(\psi, t) = \frac{v_m}{k_m} \begin{cases} 1 - \frac{\sinh \frac{L(t)-w(t)}{\lambda}}{\sinh \frac{L(t)}{\lambda}} \cosh \frac{\psi(t)}{\lambda} & \psi(t) < w(t), \\ \frac{\sinh \frac{w(t)}{\lambda}}{\sinh \frac{L(t)}{\lambda}} \cosh \frac{L(t)-\psi(t)}{\lambda} & \psi(t) > w(t). \end{cases} \tag{S9}$$

In limit where $\lambda \ll L$, this simplifies to

$$\tilde{m}(t; x_0) = \frac{v_m}{k_m} \begin{cases} 1 - e^{-w(t)/\lambda} \cosh \frac{\psi(t)}{\lambda} & \psi(t) < w(t) \\ \sinh \frac{w(t)}{\lambda} e^{-\psi(t)/\lambda} & \psi(t) > w(t). \end{cases} \tag{S10}$$

#### S3. QUANTIFICATION OF PATTERN AND THE RESPECIFICATION TIME $\tau$

In order to explore patterning dynamics, we define a boundary position, $\chi$, that delineates cells at states with high $n_1$ from cells at states with high $n_2$. We use thresholding of the $n_2$ concentration profile to find the position $\chi$ when the normalised concentration of $n_2$ reaches a fraction $f$ of the maximum amplitude given by the production rate over the effective degradation rate (Fig. S1(a)),

$$n_2(\chi(t), t) = f \frac{v_2}{k_2 + g} = f \max(n_2(x, t)). \tag{S11}$$

In the case where the $n_2$ concentration is above the threshold for all $x$ or below the threshold for all $x$, no boundary exists.

In the main text, we define the respecification time $\tau$ as the time it takes a cell initially in the high $n_1$ state to transition to an $n_2$ state which we define as reaching a given threshold of $n_2$ concentration upon exiting the bistable region (Equation 6, Fig. 3, Fig. S7(b)). We employ two methods to quantify $\tau$: a full spatial solver of $m, n_1$ and $n_2$ using Equations 1, 2 and 3 in the main text; and a non-spatial solver using an analytical expression for $\tilde{m}$ input to numerically find $n_2$. The value of $\tau$ obtained from the spatial solver agrees well with the value obtained from the non-spatial solver with analytically calculated $\tilde{m}$ (Fig. S7(c)).

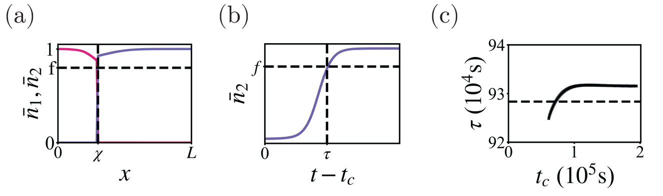


FIG. S1. (a) Normalised gene $\bar{n}_1$ and $\bar{n}_2$ concentration over space $x$. The boundary position $\chi$ is defined as the position where $n_2$ reaches a fraction $f$ of its maximum $v_2/(k_2+g)$. (b) Normalised gene concentration $\bar{n}_2$ against time since leaving the bistable region, $t - t_c$. (c) The respecification time $\tau$ for cells leaving the bistable region at time $t_c$, quantified in the 1D spatial simulation for cells as they leave the bistable region at time $t_c$ (circles) compared to a non-spatial approximation from an analytical input of $\tilde{m}$ (dashed line).

#### S4. PATTERN SCALING IN A MONOSTABLE REGIME

In the main text we find scaling can emerge from the non-negligible relaxation times of the GRN in response to changes in the morphogen signal experienced by a cell as it is advected away from the morphogen source in a growing tissue, $\tilde{m}$. The distance over which the system scales due to the GRN dynamics, which we term $\Delta$, depends on the relative timescales of advection and GRN processing. This scaling emerges independent of multistability, therefore we predict that scaling will emerge even when the system is in a monostable regime. In this section, we simulate the toggle switch in a monostable regime and quantify scaling in a growing tissue.

The stability of the morphogen-induced toggle switch depends on the Hill coefficients $p, q$ and the amplitudes of the genes $v_1/k_1$ and $v_2/k_2$. A simple choice of parametrisation that yields a monostable regime is setting the Hill coefficients to unity, $p = q = 1$. This means there are no bifurcations in response to changes in $m$, and there is only a single steady state solution for $n_1$ and $n_2$ for a given input of $m$ across space (Fig. S2(a,b)). For $p = q = 1$, solutions for $n_1$ and $n_2$ at steady state, $n_1^*$ and $n_2^*$ can be found analytically (Fig. S2(c)),

$$n_1^* = \frac{1}{2}\Bigg( -1 + \frac{v_1 m^h}{k_1(1+m^h)} - \frac{v_2}{k_2} + \sqrt{\left(1 - \frac{v_1 m^h}{k_1(1+m^h)} + \frac{v_2}{k_2}\right)^2 + 4\frac{v_1 m^h}{k_1(1+m^h)}}\Bigg) \tag{S12}$$

$$n_2^* = \frac{1}{2}\Bigg( -1 - \frac{v_1 m^h}{k_1(1+m^h)} + \frac{v_2}{k_2} + \sqrt{\left(1 + \frac{v_2}{k_2} - \frac{v_1 m^h}{k_1(1+m^h)}\right)^2 + 4\frac{v_2}{k_2}}\Bigg) \tag{S13}$$

In a monostable regime, there is a well-defined morphogen value at which the gene profiles reach a given threshold concentration at steady state, that can be read off the $n_1^*$ and $n_2^*$ against $m$ graphs (Fig. S2(a,b)). The position at which this $m$ value is met is the steady state $n_1$ versus $n_2$ boundary position (Fig. S2(c)). Unlike in the bistable case, the steady state boundary position is independent of initial conditions.

To test if scaling can emerge in the absence of bistability in growing tissues, we next simulate a growing tissue. When we allow the system to grow, the boundary position reaches a higher value than in the static case (Fig. S2(d)). The maximum value of the boundary position, which we term $\chi^*$, increases with the growth rate, (Fig. S2(e)), in a similar manner to the bistable case once the boundary position has surpassed the edge of the bistable region (Fig. 2(d) in the main text).

The analysis presented in the main text shows that scaling emerges in the toggle switch from the relaxation of the GRN occurring over a given timescale while cells are being advected through the tissue. The same analysis can be performed in the monostable case, by defining $\tau$ as the time it takes to transition from $n_1$ to $n_2$ state once cells move out of the $n_1$ region. As with the bistable case, we can quantify $\tau$ and input into Equation in the main text, and we find this matches well with the numerical simulations, confirming that scaling due to GRN relaxation can occur in the absence of multistability (Fig. S2(e)). Furthermore, for a monostable system, scaling is completely absent in the case of slow growth as predicted, and the boundary position relaxes rapidly to the steady state value matching the analytical expressions (Fig. S2(f)).

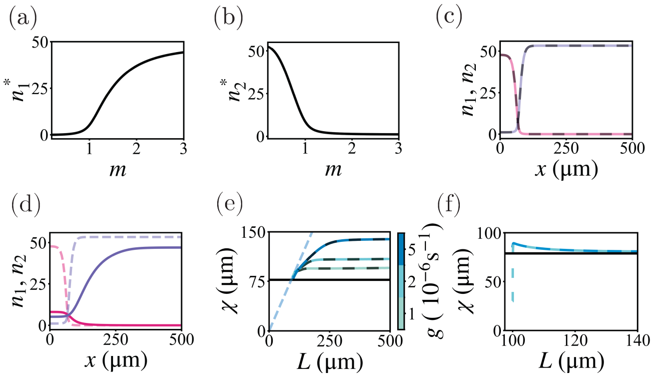


FIG. S2. Monostable system, with Hill coefficients $p = q = 1$. (a) Steady state concentration $n_1^*$ against morphogen concentration $m$. (b) Steady state concentration $n_2^*$ against morphogen concentration $m$. (c) $n_1$ (pink) and $n_2$ (purple) concentrations in space $x$ for a numerically simulated non-growing system, and the analytical solution from Equations and (black dashed line), (d) $n_1$ (pink) and $n_2$ (purple) concentrations in space $x$ for a growing system (solid lines) compared to a non-growing system (dashed lines). (e) Position of boundary $\chi$ against length of tissue $L$. Analytical solutions from numerical quantifications of $\tau$ inputted into Equation in the main text are plotted in black dashed lines. (f) Position of boundary $\chi$ against length of tissue $L$ for a tissue at the limit of slow growth. For growing systems with a small growth rate, $\chi$ does not scale and instead relaxes to the steady state boundary position obtained from the analytical equations $\chi_0$ (black), for initial conditions of $n_1, n_2 = 0$ (green dashed) and $n_1, n_2$ initiated at steady state (blue).

#### S5. APPLICATION TO SELF-ACTIVATING NODE

To illustrate that GRN multistability and resulting capacity for hysteresis drives scaling independent of the specific wiring of networks, we demonstrate that scaling mediated by multistability can also emerge in other minimal networks. We use the case of the self-activating node where a morphogen $M$ activates the production of a node $N$, which also self-activates (Fig. S5(a)), with Hill coefficients $H$ and $P$ respectively. Analogous to the toggle switch, the dynamics of the gene concentration are described by production, degradation, dilution and advection,

$$\frac{\partial n}{\partial t} = v_n \frac{m^H}{1+m^H} \frac{1}{1+n^P} - (k_n + g)n - u_x \frac{\partial n}{\partial x}, \quad \text{(S14)}$$

where we have non-dimensionalised by the half maximum concentration for the Hill functions, as in the toggle switch case described above $m = M/M^*, n = N/N^*$, $k_n$ is the degradation rate of $n$, and $v_n$ is the normalised production rate.

Next, we consider the bifurcation diagram for the self-activating node (Fig. S5(a)). There exists a stable solution for $n = 0$ for all values of the morphogen, and at a critical morphogen concentration, the system undergoes a saddle node bifurcation (Fig. S5(a)), meaning that bistability occurs for all morphogen values above this critical value. We thus predict that a boundary can scale up until the edge of the bistable region defined by this critical morphogen concentration.

We define a boundary as the position at which the concentration $n$ reaches a threshold value, analogous to in the toggle switch (Equation S3). To explore if scaling can emerge, we simulate a self-activating node network in a growing tissue and quantify the boundary dynamics. We find the $n$ profile and the boundary position $\chi$ scale up until a maximum value, as predicted (Fig. S5(b)). For small growth rates, the maximum boundary position is constrained by the bistable region, and as we increase the growth rate, the maximum value increases, as in the toggle switch (Fig. S5(c)).

As with the toggle switch, we can predict the maximum boundary position a system scale for a given growth rate from a numerical estimation of $\tau$, the respecification time. If we compute $\tau$ for the self-activating node and input into Equation 7 in the main text, we find this agrees well with numerical simulations, confirming that the mechanism of scaling arising from GRN dynamics applies to

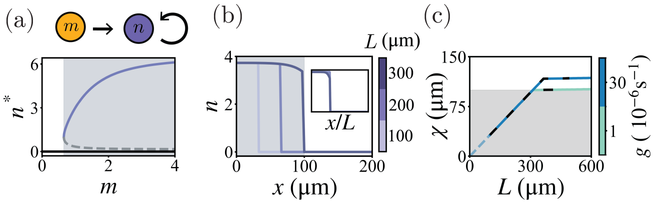

FIG. S3. (a) Graph and bifurcation diagram for a self-activating node, displaying the value of $n$ at the fixed point, $n^*$ against morphogen concentration $m$. (b) $n$ profiles for different lengths (colour bar) in absolute space $x$ and relative space $x/L$. (c) Boundary position $\chi$ against tissue length $L$ for different growth rates $g$ (colour bar), numerical solutions (solid) and solutions from numerical quantifications of $\tau$ inputted into Equation 7 in the main text (black, dashed).

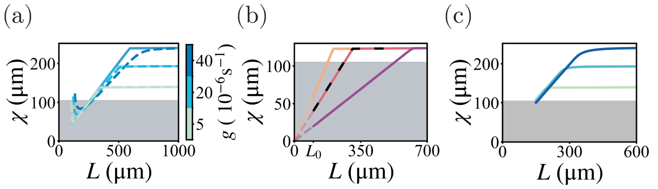

FIG. S4. The effect of initial conditions of $n_1, n_2$ and $m$. (a) Boundary position $\chi$ against tissue length $L$ for system is initiated at a steady state profile for $n_1$ and $n_2$ (solid lines) at initial length $L_0$ versus systems systems initiated at $n_1 = n_2 = 0$ for all $x$ (dashed lines), for different growth rates $g$ (colourbar). (b) Boundary position $\chi$ against tissue length $L$ for systems initiated with $n_1 = v_1/k_1, n_2 = 0$ for $x < x_b$, $n_1 = 0, n_2 = v_2/k_2$, elsewhere, for $x_b = 0.2$ (purple), $x_b = 0.4$ (red), and $x_b = 0.6$ (orange), and when $n_1, n_2$ are initiated at steady state for a static tissue (black, dashed). (c) Boundary position $\chi$ against tissue length $L$ for a system initiated with a boundary outside of the bistable region for different growth rates $g$ (colourbar in (a))

the self-activating node. This confirms our understanding that the scaling properties emerge due to multistability and is independent of the underlying wiring that gives rise to it.

#### S6. INITIAL CONDITIONS

In this section, we examine the effects of the initial conditions for the morphogen and genes on the resulting boundary dynamics and emergence of scaling.

All cases in the main text are initiated at steady state profiles of $m, n_1$ and $n_2$ evaluated at a static length $L_0$, with an effective degradation rate for the genes equal to $g+k_1$ and $g+k_2$. When we initiate the system away from steady state, we find that the resulting dynamics depend on the relative growth rate compared to the GRN dynamics. For small growth rates compared to the GRN dynamics, initiating the system at $n_1 = n_2 = 0$ for all $x$ has a minor effect on the early dynamics. The system approaches the steady state of $n_1$ and $n_2$ on approximately the timescales determined by the gene degradation rates $1/k_1, 1/k_2$, which is fast compared to growth. The boundary at tissue lengths close to the initial length is similar to the steady state boundary position, and the boundary dynamics collapse on to the case where the system is initiated at steady state (Fig. S4(a)). For larger growth rates, $g > k_1, k_2$, the timescales $1/k_1$ and $1/k_2$ are no longer faster than the timescale of growth $1/g$, and so the build up dynamics occur over a larger span of tissue lengths and disrupt scaling over the build up timescale dictated by the GRN dynamics. Once the boundary has been established and is close to the steady state value, scaling is reinstated (Fig. S4(a)).

We next consider the effects of initiating the system with step-like $n_1$ and $n_2$ profiles with a boundary within the bistable region. A boundary can be initiated at any position within the bistable region and scaling is observed until a maximum boundary position value $\chi^*$ independent of initial conditions (Fig. S4(b)). This results in a different pattern proportion $\chi_0/L_0$ which dictates the factor of proportionality between $\chi$ and $L$. In addition, the different values of the initial boundary affects the advection velocity of the $n_1$ cells at the boundary. It follows that, the position of the initial boundary impacts the range of tissue lengths over which a system scales (Fig. S4(b)). Finally, initiating the system at steady state, we are simply allowing the GRN parameters to determine initial boundary position to be scaled (Fig. S4(b)).

In the main text, we find that the boundary can extend by a distance $\Delta$ outside the bistable region. We now consider initiating the system with step-like $n_1$ and $n_2$ profiles with a boundary just outside of the bistable region. We find that the boundary position will scale from the edge of the bistable region until it reaches $\chi^* = x_c + \Delta$ (Fig. S4(c)).

#### S7. EFFECT OF FIXED POINT DECAY ON BOUNDARY DYNAMICS

In section 3, we define the boundary position as the position where the $n_2$ concentration profile meets a threshold, and we use this quantification to explore pattern scaling. We choose to use $n_2$ to define the boundary rather than $n_1$ because the amplitude of $n_1^*$, the value of $n_1$ at the fixed point, is directly modulated by the morphogen concentration, causing the magnitude of the $n_1$ fixed point to decay across the tissue. In this section, we discuss the effect of $n_1$ fixed point decay on boundary quantification.

The bifurcation diagram of $n_2^*$ against $m$ in the main text (Fig. 1(b)) shows that the values of $n_2^*$ at the high $n_2$ fixed point is very close to $v_2/k_2$ across the bistable region. The respective bifurcation diagram of $n_1^*$ against $m$, however, shows that, for some parameter choices, the magnitude of $n_1$ at the fixed point can decrease by an order of magnitude across the bistable region (Fig. S5(a)).

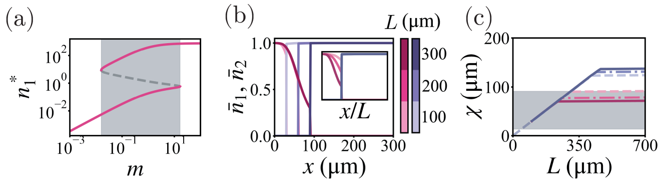


FIG. S5. (a) A bifurcation diagram showing $n_1^*$, the values of $n_1$ at the fixed points, against the morphogen concentration $m$. The bistable region (shaded) has 2 stable steady states (pink) and an unstable steady state (grey, dashed). (b) Normalised $n_1$ and $n_2$ concentration, $\bar{n}_1, \bar{n}_2$ against position $x$ in the case of a growing tissue, for tissue lengths $L = [100, 200, 300]\mu$m (colourbars). Insets show the same profiles in relative space $x/L$. (c) Boundary position $\chi$ against tissue length $L$ for different values of $f$ for calculating the boundary position, for a threshold $fv_i/(k+g)$, where $i = 1$ for $n_1$ (pink) and $i = 2$ for $n_2$ (purple). $f = [0.2, 0.5, 0.8]$, for dashed, dot-dash and solid lines respectively.

In a growing tissue, cells may stay at the $n_1$ fixed point as they are advected across the tissue, but the position of the $n_1$ fixed point moves in $n_1 - n_2$ space. This affects scaling of the $n_1$ profile (Fig. S5(b)).

The decay of the $n_1$ fixed point leads to a less sharp boundary, which means that the fraction $f$ chosen to quantify the boundary affects the position more so than in the case where the boundaries are sharp (Fig. S5(c)). This can lead to the threshold position where $n_1 = fv_1/(k_1 + g)$ being inside the bistable region, and lead to gap between the positions of the $n_1$ and $n_2$ boundaries.

#### S8. BOUNDARY DYNAMICS AND $\chi^*$

In this section, we show the non-normalised profiles of $n_1$ and $n_2$ as presented in Fig. 2(c) of the main text, show the details of how we varied the position of the bistable region in Fig. 2(e) on the gene expression profiles and demonstrate the agreement between the analytical equations and numerical simulations of the boundary position dynamics.

In the main text, we demonstrate scaling of $n_1$ and $n_2$ profiles in a growing tissue and show normalised profiles $\bar{n}_1, \bar{n}_2$, where

$$\bar{n}_1 = \frac{n_1(k_1 + g)}{v_1}, \tag{S15}$$

$$\bar{n}_2 = \frac{n_2(k_2 + g)}{v_2}. \tag{S16}$$

For completeness, we include the non-normalised $n_1$ and $n_2$ profiles in a growing tissue (Fig. S6(a)), corresponding to the same system as the normalised $n_1$ and $n_2$ in the main text (Fig. 2(c)). Scaling emerges regardless of normalisation.

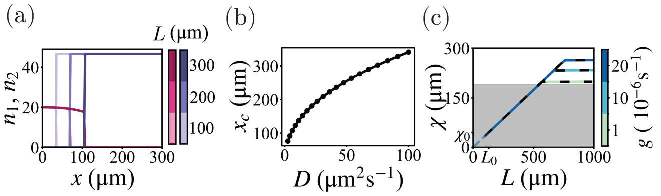


FIG. S6. (a) $n_1$ and $n_2$ concentration profiles for different tissue lengths (colourbar) for a growing tissue. (b) Position of the edge of the bistable region $x_c$ against morphogen diffusion coefficient $D$. (c) Boundary position $\chi$ against tissue length $L$ for growing tissues, with different growth rates (colourbar) from numerical solutions (solid, light to dark blue) and analytical solutions (dashed, black) obtained from Equation S8. Pattern boundary extends beyond the bistable region (grey, shaded).

In the main text, we demonstrate that scaling emerges up until a maximum boundary position, $\chi^*$. In Equation S2 we find an expression for $\chi^*$ as a function of the edge of the bistable position $x_c$ and $\Delta$, which is a function of the respecification time $\tau$. To explore the relationship between the edge of the bistable region $x_c$ and the maximum boundary position $\chi^*$, we vary the edge of the bistable region and find the effect on $\chi^*$ (Fig. 2(e) in the main text). The mechanism by which we varied $x_c$ was by changing the underlying morphogen gradient by varying $D$, the morphogen diffusion coefficient. This affects the position at which $m = m_c$ (Fig. S6(b)).

Finally, in the main text, we present a physical rationale for how the maximum boundary position depends on the bistable region and GRN dynamics. This allows us to write the full boundary dynamics over time,

$$\chi = \begin{cases} \chi_0 e^{gt/(1+\varepsilon)} & t < t^* \\ \chi^* & t > t^*, \end{cases} \tag{S17}$$

where $t^*$ corresponds to the time where $\chi_0 e^{gt/(1+\varepsilon)} = \chi^*$. This equation arises from considering evolution of the position $\psi(t; \chi_0)$ over time for a cell that starts at the boundary $\chi_0$. Given that growth is spatially uniform, we can write this expression in terms of tissue length $L$,

$$\chi = \begin{cases} \frac{\chi_0 L}{L_0} & L < L^* \\ \chi^* & L > L^*, \end{cases} \tag{S18}$$

where $L^*$ is the tissue length where $\chi_0 L/L_0$ reaches $\chi^*$. This expression demonstrates the linear scaling that emerges for $L < L^*$. Using a numerical input of $\tau$, we calculate the value of $\chi^*$, and plot Equation S8 and find it in good agreement with the numerical simulations (Fig. S6(c)).

#### S9. DEPENDENCE OF $\tau$ ON PARAMETERS

In the main text we have examined the dependence of the timescale of GRN respecification $\tau$ on $k_2$, the degradation rate of $n_2$ (Fig. 3d), and $g$, the growth rate (Fig. 3g). In this section, we provide additional intuition of the relationships between $\tau$ and the GRN parameters, and consider how $\tau$ changes in response to changes in $k_1$, the degradation rate of $n_1$;, $v_2$, the production rate of $n_2$; and to changes in both gene degradation rates concomitantly.

What determines the tempo of $n_1$ and $n_2$ dynamics is complex in non-linear systems. The rate of change in genes depends on the intrinsic rates of the system $(k_1, k_2, v_1, v_2)$, the position in $n_1 - n_2$ space, and the current $m$ value. In the case of $\tau$, we narrow our analysis of tempos down to focus on the timescale of gene dynamics just after the saddle-node bifurcation (Fig. S7(b)). By definition, upon exit of the bistable region, the high $n_1$ fixed point disappears, however, in the vicinity of the saddle-node bifurcation there exists a ghost of the fixed point. This ghost slows the GRN dynamics in a phenomenon described as critical slowing down [4]. In our system, the effect of critical slowing down depends how persistent the ghost is as $\tilde{m}$ decreases and on the rate of advection just beyond the bistable region which determines $\tilde{m}$. In the limit of fast advection, where the morphogen concentration experienced by a cell immediately decays to $\tilde{m} = 0$ upon exiting the bistable region, there is no production of $n_1$, and thus $\tau$ is limited only by how quickly the current $n_1$ concentration can decrease and $n_2$ can increase. This corresponds to the fast advection case presented in the main text (Fig. 3(g)). On the contrary, for slow advection, the scale of $\tau$ is dominated by how slow the GRN dynamics are in the vicinity of the bifurcation. In summary, the time of respecification is controlled by a combination of GRN speed and advection velocity, both evaluated by the bistable region.

These two modes of action on the magnitude of $\tau$ can be changed independently, for example by modulating the GRN parameters, or changing the morphogen profile to move the bistable region. Growth, however, affects both GRN speed and advection. We note in the main text that growth has antagonistic effects on the extension to scaling: increasing the rate at which the system moves away from the ghost and increasing dilution will speed up the GRN dynamics and decrease $\tau$, which in isolation would lead to a smaller $\Delta$. However, a larger advection velocity means the system travels further in the same time. In general, increasing growth leads to a larger $\Delta$, which can be rationalised by differentiating the expression for $\chi^*$ and noting that increasing growth increases $g$ linearly but decreases $\tau$ sublinearly (Fig. 3(d) in the main text). This results in the exponent $g\tau(g)/(1+\varepsilon)$ increasing with increasing $g$.

Finally, we consider the effect on $\tau$ of other rates present in the GRN formulation. Increasing the degradation rate $k_1$ causes a decrease in $\tau$ that is more non-linear than in response to decrease in $k_2$ (Fig. S7(a)). This represents how the decrease in $n_1$ is the limiting step in moving from the high $n_1$ ghost toward the high $n_2$ fixed point. The quantitative difference in $\partial\tau/\partial k_2$ compared to $\partial\tau/\partial k_1$ could be inferred from how the velocities $dn_1/dt$ and $dn_2/dt$ vary in magnitude close to the ghost. The most rapid decline in $\tau$ in response to change in $k_1$ occurs when $k_1 \approx k_2$. As $k_1$ approaches higher values, $\tau$ plateaus as it ceases to become the limiting rate. In the symmetrical case $k_1 = k_2 = k_i$, $\tau$ decreases rapidly in response to decrease in $k_i$ (Fig. S7(b)). When $k_i$ reaches too large a value, the system is pushed out of the bistable regime. The definition of $\tau$ from the time a cell exits the bistable region is therefore inapplicable. We instead find the characteristic timescale for a cell to become the boundary after passing the value of the boundary with no growth, as defined for the monostable case discussed above. Increasing the production rate $v_2$ leads to a decrease in $\tau$ (Fig. S7(c)). The effect is less significant over several orders of magnitude than the degradation rates, indicating that the production rates are less critical in dictating $\tau$.

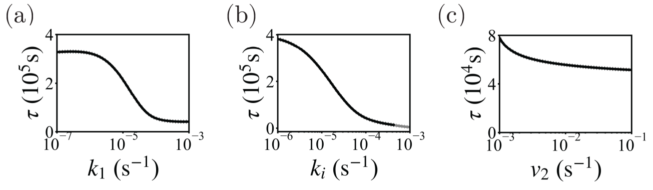


FIG. S7. Quantification of $\tau$ for the same parameters as Fig. 3 in the main text. (a) Increasing the degradation rate $k_1$ decreases $\tau$ in a non-linear fashion. (b) Varying both $k_i = k_1 = k_2$ together similarly decreases $\tau$. For large values of $k_i$, there is no bistable region, and thus $\tau$ is measured as the time to respecify after a system cross the morphogen concentration corresponding to the boundary position in the no growth case, as in the monostable case. (c) Increasing the production rate of $n_2$, $v_2$ leads to a modest decrease of $\tau$.

#### S10. SOURCE GROWTH

In the main text, we explore how growth of the morphogen source accompanying tissue growth can affect scaling of systems patterned with GRNs. We find that source growth translates the bistable region as the morphogen profile evolves, expanding scaling. In this section, we provide derivations for the results concerning the translation of the bistable region and the optimal values of $\gamma$ for linear scaling. Finally, we provide details for the boundary dynamics in the regime following exit from the bistable region.

In the main text, we consider how source growth affects boundary position dynamics by moving the edge of the bistable region. We can understand analytically how source growth affects the edge of the bistable region by considering where the threshold $x_c$ evolves as a function of the tissue size. If we assume the edge of the bistable region is outside of the source itself and consider the limit

$\lambda \ll L$ for simplicity, Equation S8 yields

$$x_c(L) = \lambda \ln \left[ \frac{v_m \sinh\left(\frac{\beta L^{\gamma})}{\lambda}\right)}{m_c k_m} \right]. \quad \text{(S19)}$$

This expression allows us to analytically find the dynamics of $\chi$ for time $t > t^*$, Equation 8 in the main text. The time $t^*$ is the time at which a boundary ceases to scale at the advection rate and instead traces the evolution of the bistable region, and occurs when the boundary exits the bistable region plus the additional distance $\Delta$. We find $t^*$ by equating a linearly scaling boundary with the edge of the bistable region as discussed in the main text. In general, the boundary may exit the bistable region from the upper edge, $x_c$, as presented in the main text, or from the lower edge $x'_c$, depending on the relative rates of advection of the boundary compared to the translation of the bistable region. For weakly non-linear source growth where $\gamma \approx 1$, it is unlikely for the source to be growing fast enough to overtake advection of the boundary, hence we focus on the case where the boundary reaches the upper edge.

In the main text, we state that constraints on $\gamma$ can be found that increase the values of $\chi_t$ and $t^*$, to extend the period over which the system scales at the advection rate by delaying the point at which the boundary exits the bistable region (Fig. 4(f) in the main text). These constraints can be found from considering how $\gamma$ affects the time $t^*$ and corresponding tissue length $L^*$ at which the boundary position reaches the edge of the bistable region, which we detail here.

Equation S19 can be rearranged to find the value of $\gamma$ such that a boundary scaling due to advection in the bistable region reaches the edge of the bistable region given by $m = m_c$ at tissue length $L^*$,

$$\gamma^b(m_c) = \frac{\ln(\frac{\lambda}{\beta} \sinh^{-1}(\frac{k_m m_c}{v_m} e^{\chi_0 L/L_0 \lambda}))}{\ln(L^*)}. \quad \text{(S20)}$$

For the boundary position to be within the bistable region at tissue length $L^*$, $\gamma$ must be within the bounds on $\gamma$ corresponding to the upper and lower edges of the bistable region,

$$\gamma^b(m_c) < \gamma^* < \gamma^b(m'_c), \quad \text{(S21)}$$

However, this does not preclude the boundary from exiting the bistable region at an earlier tissue length. In order to scale over a given length span $[L_0, L_f]$ $\gamma$ must stay within the corresponding bounds for all $L$ within $[L_0, L_f]$. Therefore, for a boundary to scale from $[L_0, L_f]$, the value of $\gamma$ must exist within a more constrained window,

$$\max[\gamma^b(L)] < \gamma^* < \min[\gamma^{b'}(L)] \quad \forall L \in [L_0, L_f] \quad \text{(S22)}$$

A value $\gamma^*$ exists for $\max[\gamma^b(L)] < \min[\gamma^{b'}(L)]$. Plotting Equation S20 over $L$ for $m_c$ and $m'_c$ shows that the bounds on $\gamma$ over time become more narrow, Fig. S8(a). The range within which $\gamma^*$ can exist scales as $\delta\gamma \sim 1/L^* \ln L^*$.

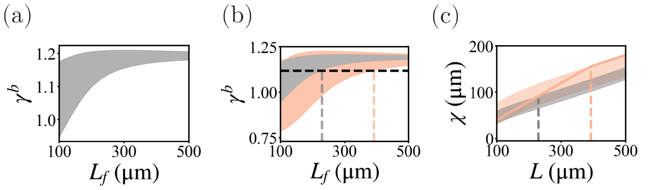


FIG. S8. (a) Bounds on $\gamma$, $\gamma_b$ in order for a linearly scaled boundary to be within the bistable region at a tissue length $L^*$ (shaded). (b) Bounds $\gamma_b$ for a slow-growing case where $\Delta = 0$, (grey, shaded) and for a faster growing case with non-zero $\Delta$ (orange, shaded). For a given $\gamma = 1.12$ (black dashed line), the boundary position will cease to linearly scale at a larger tissue size for the faster growing case (orange dashed line) compared to the slow growing case (grey dashed line). (c) Boundary position $\chi$ against tissue length $L$ for a fast growing case (orange). A slow growing case would exit the bistable region (shaded grey) earlier (grey dashed line) than the fast growing case (orange dashed line) as growth extends the scaling region by $\Delta$ (shaded orange).

As in the static source case, the edge of the bistable region imposes a limit on how far cells can carry a boundary. The limits imposed can be stringent: in the example plotted, for most values of $\gamma$, the boundary exits the bistable region after only a short amount of growth, Fig. S8(a). In the static source case, the edge of the bistable region imposes a static limit on how far cells can carry a boundary, and we found that increasing the growth rate extends the period of scaling, effectively expanding the bistable region by a distance $\Delta$. We now examine the effect of increasing growth rate to the linear scaling limits on $\gamma$.

$\Delta$ can relax the limits on $\gamma$ (Fig. S8(b)). Akin to the case of a static source, a faster growth rate allows the boundary to be maintained at positions outside of the bistable region due to the slow relative reaction time of the GRN to the new signalling environment. The slow GRN dynamics compared to growth effectively extends the bistable region, and creates a buffer zone that allows the boundary position to linearly scale over a longer period compared to the slow-growing case (Fig. S8(c)). Increasing the growth rate can relax the limits on $\gamma$ by introducing a buffer zone, Fig. S8(b). This allows the boundary position to scale at the advection rate over a longer period and expand the bounds on $\gamma^*$ compared to the slow-growing case (Fig. S8(b)). The effects of these on pattern scaling is that a faster growing system can scale for longer for a given source scaling than a slow growing system, or for a system without source scaling.

In the main text, we adapt the boundary position dynamics to account for a time dependence of the bistable region $x_c$ (Equation 8 in the main text). Once the boundary position has reached $\chi_t$, the maximum limit of the bistable region plus its extension, its evolution depends

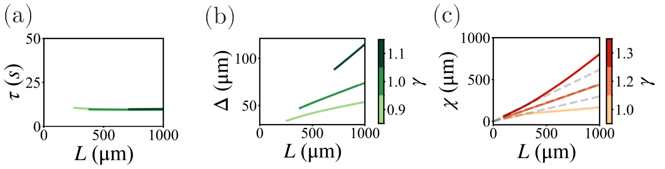


FIG. S9. (a) $\tau$ over $L$ for different values of $\gamma$ (green colourbar) shows no significant change in $\tau$. (b) $\Delta$ changes over $L$ as the edge of the bistable region extends further into the tissue, and thus advection at the edge of the bistable region increases, despite $\tau$ staying approximately constant. (c) Limited scaling can be achieved without bistability or $\Delta$, through superlinear source growth achieved by varying $\gamma$ (red colourbar) for a short span.

on the evolution of the bistable region,

$$\chi^* = x_c + \Delta = x_c(t-\tau)e^{g\tau/(1+\epsilon)}. \tag{S23}$$

Considering Equation S10, it is possible that Equation S2 may have other time dependencies: in general, $\tau$ may be time dependent, and thus may affect the dynamics of $\chi$. In the regimes examined, however, we find that $\tau$ stays constant over time, Fig. S9(a). Even if $\tau$ remains constant, $\Delta$ may increase in time, as the advection velocity increases with $x_c$, however, we find the changes in $\Delta$ to be much smaller in magnitude compared to the evolution of $x_c$, Fig. S9(b). This can be understood from differentiating $\Delta$ with respect to $x_c$: any change in $\Delta$ depends only on the change of $x_c$ between a time $t$ and $t+\tau$, which is small.

Given that $\tau$ and $\Delta$ do not vary over time, the dynamics of $\chi$ at later times is dominated by the evolution of the bistable region $x_c$. We compare these dynamics to slow-growing, monostable cases, where there are no effects of GRN dynamics, and the boundary position occurs at a given threshold morphogen concentration. In simulations of monostable systems at the limit of small growth, the boundary position increases due to source growth as the position of a morphogen concentration threshold is pushed into the tissue Fig. S9(c). For weakly superlinear source growth, approximately linear pattern scaling can be achieved without multistability or $\Delta$ for a limited period of growth, Fig. S9(c).

#### S11. PARAMETERS

In this section, we detail the parameters used for the simulated data used to generate figures throughout the article.

As discussed in the main text, we consider a scenario where the morphogen gradient is close to steady state while the GRN dynamics and growth dynamics are ongoing and focus on the regime where the morphogen degradation rate $k_m$ is larger than the degradation rates $k_1$ and $k_2$ and the growth rate $g$, $k_m \gg k_1, k_2, g$.

We choose the morphogen degradation rate $k_m$ to be on the order of $1\text{h}^{-1} = 10^{-4}\text{s}^{-1}$ and the morphogen diffusion coefficient $D$ on the order $10^{-1}\mu\text{m}^2\text{s}^{-1}$ [5], consistent with estimates of morphogens such as Shh in the mammalian neural tube [6], Dpp in the *Drosophila* wing disc [7], and Nodal and Lefty in zebrafish embryogenesis [8] .

For dorsoventral patterning in the developing mouse neural tube, the growth rate $g$ is on the order of $10^{-5}\text{s}^{-1}$, and the anisotropy parameter $\varepsilon = 0.6$, and the initial length $L_0 = 10^2\mu\text{m}$ with approximate a 5 fold change in length [9]. The anisotropy parameter in the *Drosophila* wing disc is estimated to be around $\varepsilon = 0.8$, and the growth rate on the order of $10^{-5}\text{s}^{-1}$ [2, 10]. We consider length changes of between 3 fold and 10 fold. We set the source width to be around $10^1 - 10^2\mu\text{m}$ for the static source, and choose the source ratio parameter $\beta$ such that the source ratio $w/L$ is approximately 10% [10, 11].

For the gene regulatory network parameters, we choose degradation rates on the same or similar order as the growth rate to explore cases where growth and GRN dynamics are operating at similar timescales [6]. We choose Hill coefficients $> 1$ to access the non-linear properties of GRNs, but do not choose coefficients above 3 ($p, q \leq 3$ for the results presented in this paper).


[1] P. Fried and D. Iber, Nat. Commun. **5**, 1 (2014).
[2] D. Aguilar-Hidalgo, S. Werner, O. Wartlick, M. González-Gaitán, B. M. Friedrich, and F. Jülicher, Phys. Rev. Lett. **120**, 198102 (2018).
[3] X. Richard, B. Richard, C. Mazza, and J. R. van der Meer, arXiv preprint arXiv:2304.02936 (2023).
[4] S. H. Strogatz, *Nonlinear dynamics and chaos: with applications to physics, biology, chemistry, and engineering* (Chapman and Hall/CRC, 2024).
[5] K. S. Stapornwongkul and J.-P. Vincent, Nat. Rev. Genet. **22**, 393 (2021).
[6] D. Benzinger and J. Briscoe, Dev. Cell **60**, 3421 (2025).
[7] A. Kicheva, P. Pantazis, T. Bollenbach, Y. Kalaidzidis, T. Bittig, F. Julicher, and M. Gonzalez-Gaitan, Science **315**, 521 (2007).
[8] P. Müller, K. W. Rogers, B. M. Jordan, J. S. Lee, D. Robson, S. Ramanathan, and A. F. Schier, Science **336**, 721 (2012).
[9] A. Kicheva, T. Bollenbach, A. Ribeiro, H. P. Valle, R. Lovell-Badge, V. Episkopou, and J. Briscoe, Science **345**, 1254927 (2014).
[10] O. Wartlick, P. Mumcu, A. Kicheva, T. Bittig, C. Seum, F. Jülicher, and M. Gonzalez-Gaitan, Science **331**, 1154 (2011).

[11] R. D. Ho, K. Kishi, M. Majka, A. Kicheva, and M. Zagorski, PLoS Comput. Biol. **20**, e1012508 (2024).

| Symbol | Quantity | Unit | Fig. 2 | Fig. 3 | Fig. 4 |
|---|---|---|---|---|---|
| $D$ | Morphogen diffusion coefficient | $\mu m^2 s^{-1}$ | 0.25 | 0.20 | 0.20 |
| $k_m$ | Morphogen degradation rate | $s^{-1}$ | $3.5 \times 10^{-4}$ | $3.5 \times 10^{-4}$ | $4.0 \times 10^{-4}$ |
| $v_m$ | Morphogen production rate | $s^{-1}$ | 0.060 | 0.006 | 0.0023 |
| $w$ | (Initial) source width | μm | 20.0 | 10.0 | 10.0 |
| $\beta$ | Source width ratio | $\mu m^{1-\gamma}$ | - | - | 0.1 |
| $\varepsilon$ | Anisotropy parameter | - | 0.80 | 0.8 | 0.8 |
| $v_1$ | Production rate of $n_1$ | $s^{-1}$ | $6.06 \times 10^{-4}$ | 0.043 | 0.010 |
| $v_2$ | Production rate of $n_2$ | $s^{-1}$ | $4.42 \times 10^{-3}$ | $9.3 \times 10^{-4}$ | $9.3 \times 10^{-4}$ |
| $k_1$ | Degradation rate of $n_1$ | $s^{-1}$ | $1.0 \times 10^{-5}$ | $5.0 \times 10^{-5}$ | $5.0 \times 10^{-5}$ |
| $k_2$ | Degradation rate of $n_2$ | $s^{-1}$ | $7.5 \times 10^{-5}$ | $5.0 \times 10^{-5}$ | $5.0 \times 10^{-5}$ |
| $q$ | Hill coefficient for $n_2$ on $n_1$ | - | 3 | 2 | 2 |
| $p$ | Hill coefficient for $n_1$ on $n_2$ | - | 2 | 2 | 2 |
| $h$ | Hill coefficient for $m$ on $n_1$ | - | 1 | 3 | 3 |
| $g$ | Growth rate | $s^{-1}$ | $2.0 \times 10^{-5}$ | $1.0 \times 10^{-5}$ | $1.0 \times 10^{-5}$ |
| $f$ | Threshold fraction | - | 0.6 | 0.8 | 0.8 |
| $L_0$ | Initial tissue length | μm | 100 | 100 | 100 |

TABLE S1. Parameter values used for Figures 2–4 in the main text.

| Symbol | Quantity | Unit | Fig. S2 | Fig. S3 | Fig. S4 | Fig. S5 |
|---|---|---|---|---|---|---|
| $D$ | Morphogen diffusion coefficient | $\mu m^2 s^{-1}$ | 0.25 | 0.25 | 0.25 | 0.25 |
| $k_m$ | Morphogen degradation rate | $s^{-1}$ | $4.0 \times 10^{-4}$ | $3.5 \times 10^{-4}$ | $3.5 \times 10^{-4}$ | $3.5 \times 10^{-4}$ |
| $v_m$ | Morphogen production rate | $s^{-1}$ | 0.015 | 0.006 | 0.006 | 0.006 |
| $w$ | (Initial) source width | μm | 10 | 20 | 20 | 20 |
| $\beta$ | Source width ratio | $\mu m^{1-\gamma}$ | - | - | - | - |
| $\varepsilon$ | Anisotropy parameter | - | 0.8 | 0.8 | 0.8 | 0.8 |
| $v_1$ | Production rate of $n_1$ | $s^{-1}$ | 0.001 | - | $1.61 \times 10^{-3}$ | $6.1 \times 10^{-4}$ |
| $v_2$ | Production rate of $n_2$ | $s^{-1}$ | 0.004 | - | $1.25 \times 10^{-3}$ | $4.4 \times 10^{-3}$ |
| $k_1$ | Degradation rate of $n_1$ | $s^{-1}$ | $1.0 \times 10^{-5}$ | - | $7.0 \times 10^{-5}$ | $1.0 \times 10^{-5}$ |
| $k_2$ | Degradation rate of $n_2$ | $s^{-1}$ | $7.5 \times 10^{-5}$ | - | $4.0 \times 10^{-5}$ | $7.5 \times 10^{-5}$ |
| $v_n$ | Production rate of $n$ | $s^{-1}$ | - | $1.0 \times 10^{-4}$ | - | - |
| $k_n$ | Degradation rate of $n$ | $s^{-1}$ | - | $1.5 \times 10^{-5}$ | - | - |
| $q$ | Hill coefficient for $n_2$ on $n_1$ | - | 1 | - | 3 | 3 |
| $p$ | Hill coefficient for $n_1$ on $n_2$ | - | 1 | 2 | 2 | 2 |
| P | Hill coefficient for $n$ on itself | - | - | 2 | - | - |
| $h$ (or H) | Hill coefficient for $m$ on gene | - | 3 | 2 | 1 | 1 |
| $g$ | Growth rate | $s^{-1}$ | $5.0 \times 10^{-9}$ | $1.0 \times 10^{-5}$ | $2.0 \times 10^{-6}$ | $2.0 \times 10^{-5}$ |
| $f$ | Threshold fraction | - | 0.6 | 0.4 | 0.6 | 0.6 |
| $L_0$ | Initial tissue length | μm | 100 | 100 | 100 | 100 |

TABLE S2. Parameter values used for supplementary Figures S2–S5, S7. The parameters for Fig. S6 are the same as for Fig. 2 in the main text; for Fig. S7 the same as Fig. 3 in the main text; and Fig. S8 and S9 the same as Fig. 4 in the main text.
